# The *Pseudomonas aeruginosa* PrrF sRNAs promote biofilm formation at body temperature

**DOI:** 10.1101/2024.12.11.628005

**Authors:** Rhishita Chourashi, Jacob Weiner, Tra-My Hoang, Khady Ouattara, Amanda G. Oglesby

**Affiliations:** University of Maryland, Baltimore, School of Pharmacy, Department of Pharmaceutical Sciences Baltimore, Maryland, 21201; University of Maryland, Baltimore, School of Medicine, Department of Microbiology and Immunology, Baltimore, Maryland, 21201

**Author notes:** University of Maryland, Baltimore, School of Pharmacy, Department of Pharmaceutical Sciences Baltimore, Maryland, 21201 (Tra-My Hoang).

**Keywords:** iron regulation, temperature regulation, PrrF, type IV pili, biofilm, *Pseudomonas aeruginosa*

## Abstract

*Pseudomonas aeruginosa* is a Gram-negative opportunistic pathogen that causes both acute and chronic infections in vulnerable populations. Treatment of *P. aeruginosa* infections is increasingly challenging due to multi-drug resistance as well as biofilm formation that further increases antibiotic tolerance. Iron, which is sequestered by the host innate immune system, is also a key nutrient that required for *P. aeruginosa* biofilm formation. Previous work showed that the iron-responsive PrrF small regulatory RNAs (sRNAs), which are key to *P. aeruginosa’s* iron starvation response and required for virulence in murine lung infection, are dispensable for biofilm formation. However, these studies were performed using flow-cell biofilms at room temperature. Here we demonstrate a temperature dependency for PrrF in *P. aeruginosa* biofilm formation – the genes for these sRNAs are required for optimal biofilm formation at 37°C but not 25°C. We further show that the Δ*prrF* mutant lacks a yet-to-be identified surface appendage that is produced at 37°C but not 25°C. These studies demonstrate that the importance of the PrrF sRNAs in *P. aeruginosa* biofilm formation at body temperature and reveal a previously under-appreciated role of temperature in iron homeostasis and *P. aeruginosa* biofilm physiology.

**IMPORTANCE:** Biofilm formation is a critical aspect of pathogenesis for many microbial pathogens as it confers increased tolerance to the host immune system and antimicrobial treatments. *Pseudomonas aeruginosa* is an opportunistic pathogen that forms biofilms during infection, resulting in antimicrobial tolerance and treatment failure. Iron is a known requirement for *P. aeruginosa* biofilm formation, yet the precise role of iron homeostasis in biofilm physiology remains unclear. Here we show that temperature alters the requirement for the PrrF small regulatory RNAs, key components of *P. aeruginosa’s* iron starvation response, for biofilm formation. Specifically, PrrF is required for the optimal formation of biofilms in flow cells at 37°C but not 25°C, yet most flow-cell biofilm studies are conducted at 25°C. These results demonstrate a previously under-appreciated role of temperature in *P. aeruginosa* biofilm physiology.

## INTRODUCTION

*Pseudomonas aeruginosa* is a versatile opportunistic pathogen that causes acute lung and blood infections in cancer patients and 10% of all hospital-acquired infections (1–4). *P. aeruginosa* also causes life-long chronic lung infections in individuals with cystic fibrosis (CF) and is a significant contributor to chronic wound infections in diabetics and surgical patients (5–7). *P. aeruginosa* is innately resistant to many therapeutic agents, and the emergence of multi-drug resistant (MDR) strains of *P. aeruginosa* leads to persistent infections, longer hospital stays, and increased mortality rates (8). Biofilm formation during chronic infections further complicates treatment due to increased tolerance of these communities against antimicrobials (9). Biofilms contribute to many types of *P. aeruginosa* infections, but are most problematic in chronic infections, such as those in the lungs of CF patients. Defining the regulatory pathways that contribute to *P. aeruginosa* biofilm formation may aid in the identification of novel anti-pseudomonal therapeutics that can improve treatment outcomes for chronic *P. aeruginosa* infections.

Iron is a critical determinant of *P. aeruginosa* virulence and biofilm formation (10–12), and disrupting iron homeostasis may yield novel targets for drug development. Iron is sequestered by host proteins as part of the innate immune defense, and *P. aeruginosa* overcomes iron-limitation during infection through the expression of multiple high affinity iron uptake systems (13, 14). This includes the production of siderophores that scavenge insoluble ferric iron [Fe(III)] in aerobic environments, the uptake of ferrous iron [Fe(II)] in anaerobic environments, and the uptake and degradation of heme, a significant source of iron in the human host (15). Multiple studies have shown that disruption of either siderophore or heme uptake strongly attenuates *P. aeruginosa* virulence (10, 12, 16, 17). However, the importance of each system appears to vary amongst different infection models (reviewed by Cornelis & Dingemans (18)), with Fe(II) and heme uptake more predominant in chronic infections (19). This is likely due to reduced oxygen availability in biofilm communities, which are characteristic of chronic infections, resulting in greater ratios of Fe(II) to Fe(III) (20) and reducing the need for siderophore synthesis. To date, studies examining the role of specific iron uptake systems in *P. aeruginosa* biofilms are largely limited to the impact of siderophores (21). Specifically, the high affinity siderophore pyoverdine, which is conserved across all the pseudomonads, is required for *P. aeruginosa* biofilm formation. In contrast, pyochelin, a lower affinity siderophore produced during modest iron starvation, is not required.

*P. aeruginosa* budgets the use of iron during infection by down-regulating the expression of Fe-dependent metabolic pathways. This function is largely mediated by the PrrF small regulatory RNAs (sRNAs), which are required for virulence in an acute murine lung infection model (22). The PrrF sRNAs are produced in low iron conditions and negatively affect the levels of numerous mRNAs coding for non-essential, iron-containing metabolic proteins (23, 24). In doing so, PrrF spares the use of iron for only the most critical processes when this nutrient becomes limiting. The PrrF sRNAs are expressed by clinical isolates from acute and chronic infections (25) and detected high levels of the PrrF sRNAs in sputum isolated from CF patients, verifying they are produced during chronic CF lung infections (19). While much has been learned about the broad impact of PrrF on cell physiology and virulence, how the PrrF sRNAs contribute to survival and biofilm formation during chronic infections remains unclear.

The shift to biofilm growth during chronic infection is modulated by intracellular levels of the second messenger cyclic di-GMP (c-di-GMP) (26), the levels of which are controlled by diguanylate cyclases (DGC) and phosphodiesterases (PDE) that form and degrade c-di-GMP, respectively. Iron regulation has been linked to c-di-GMP signaling in *P. aeruginosa,* although the precise nature of this link is not known (27). Iron also promotes *P. aeruginosa* biofilm formation in a Fur-dependent manner (21), and induction of *P. aeruginosa* biofilm formation by certain antibiotics, a phenomenon that is dependent on Fe and c-di-GMP signaling (28), is dependent on the PrrF sRNAs (29). These studies demonstrate the importance of iron and PrrF in *P. aeruginosa* biofilm physiology.

Despite the accepted role of iron in *P. aeruginosa* biofilm physiology, it remains largely unknown how the PrrF sRNAs function within *P. aeruginosa* biofilms. In this vein, the work described herein challenges a conclusion that has been held in the *P. aeruginosa* biofilm community for 20 years: that the PrrF sRNAs are not required for biofilm formation (21). He we show that the PrrF sRNAs are indeed required for biofilm formation in a flow-cell system, but that there is a temperature dependency for this phenotype. Specifically, the Δ*prrF* mutant is defective for flow cell biofilm formation at 37°C (body temperature) but not at 25°C (environmental temperature). Electron microscopy revealed the presence of extracellular appendages present on wild type PAO1 and reduced on the surface of the Δ*prrF* mutant at 37°C. Moreover, these appendages were absent in cultures grown at 25°C. Altogether, these findings indicate that regulatory pathways affecting *P. aeruginosa* biofilm formation at environmental and body temperatures are distinct.

## RESULTS

### The *prrF* promoter is transcriptionally active during 37°C biofilm growth, indicating an Fe starvation response

To begin studying iron homeostasis in *P. aeruginosa* biofilms, we generated a fluorescent transcriptional reporter using the *prrF1* promoter fused to green fluorescent protein (GFP). The reporter construct was introduced at the *att* site of strain PAO1 as previously described (30). We first confirmed that the resulting reporter strain was responsive to iron by growing it in high and low iron DTSB medium and tracking fluorescence in a BioTek Synergy HT plate reader at 37°C for 24 hours (**Fig. 1A**). We next examined the reporter strain in flow-cell biofilms formed after 48 hours at 37°C. Confocal microscopy showed fluorescence throughout the area of the biofilm that was imaged (**Fig. 1B**). This shows PrrF promoter activity is upregulated in flow-cell biofilms grown at 37°C in iron starved conditions. Together, these data indicates that P*_Prrf1_*:*gfp* reporter is responsive to iron in DTSB media and flow cell biofilm cells grown at 37°C likely express the PrrF sRNAs under iron starved conditions.

**Figure 1.**
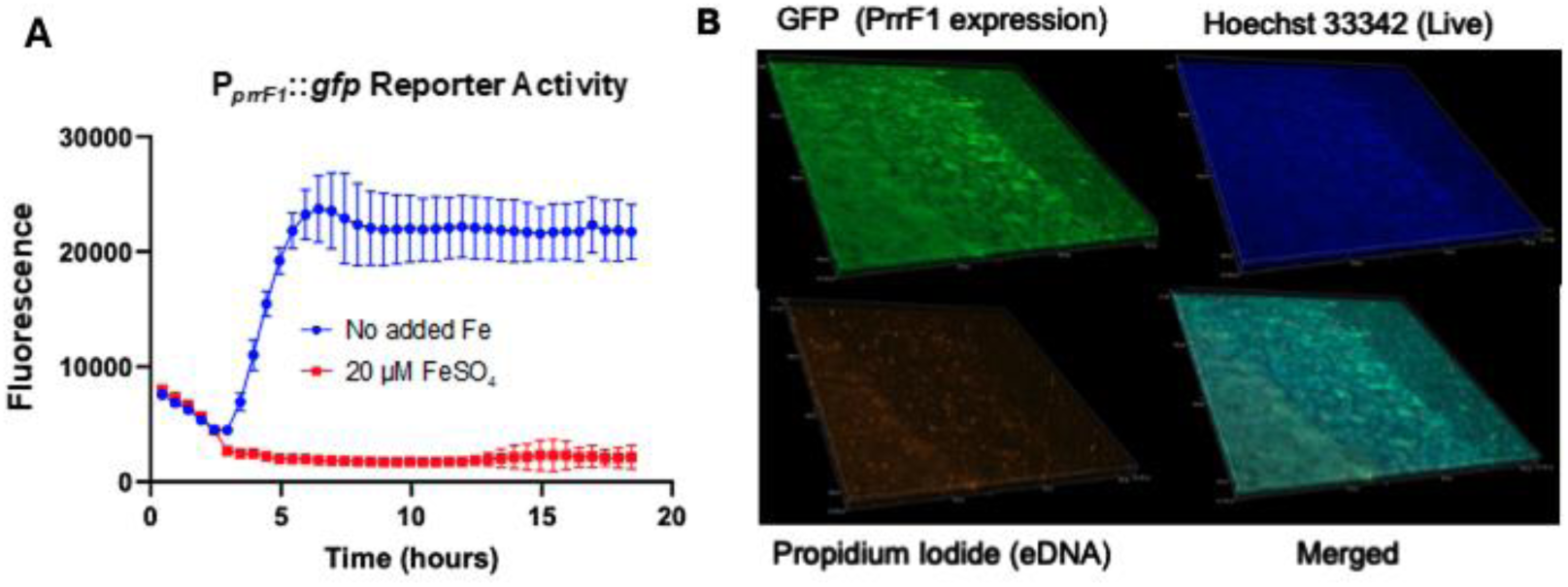
The P*_prrF1_*promoter is responsive to iron starvation in flow-cell biofilms. A. Activity in P*_prrF1_*:*gfp* reporter is responsive to iron in DTSB media. B. Representative confocal images showing PrrF1 promoter activity through GFP expression (green fluorescence) in flow-cell biofilms grown under iron limited conditions. The hoechst 33342 (blue fluorescence) stains live biofilm cells, propidium iodide (PI) stains the eDNA (extracellular DNA). The red fluorescence of PI has been pseudo-colored with orange. Images are representative of at least 5 biological replicates as shown in the Supplementary Materials (**Fig. S4**).

### The PrrF sRNAs are required for flow-cell biofilm formation at 37°C but not at 25°C

Previous studies from Banin, *et al*, demonstrated that a PAO1 Δ*prrF* mutant formed flow-cell biofilms similar to its wild-type parent (21). However, our above data indicate that the PrrF sRNAs are highly expressed in flow-cell biofilms grown at 37°C, and our earlier 37°C biofilm studies in MBEC plates suggested the Δ*prrF* mutant may exhibit altered c-di-GMP signaling in biofilms (22). Since the flow-cell biofilms from Banin and colleagues were grown at 25°C, we wondered if growth at 37°C may result in a different outcome. We therefore grew and imaged flow-cell biofilms of PAO1 and the isogenic Δ*prrF* mutant as described above. Under these conditions, the Δ*prrF* mutant demonstrated a clear defect in biofilm formation as compared to PAO1 when not supplemented with iron (**Fig. 2A-B**). Moreover, iron supplementation rescued biofilm formation of the Δ*prrF* mutant (**Fig. 2C**). Complementation of the Δ*prrF* mutant in *trans*, compared to isogenic vector-control strains, also restored biofilm formation to the Δ*prrF* mutant (**Fig. 3**). To determine if temperature is a likely reason for our findings differing from Banin, *et al*, we conducted a second series of flow-cell experiments at room temperature, imaging the biofilms at 72 hours. At this temperature, the Δ*prrF* mutant consistently formed biofilms that were either indistinguishable or slightly denser than that of the PAO1 parent strain (**Fig. 2A-B**). Combined, these data show that the PrrF sRNAs are required for flow-cell biofilm formation specifically at body temperature.

**Figure 2.**
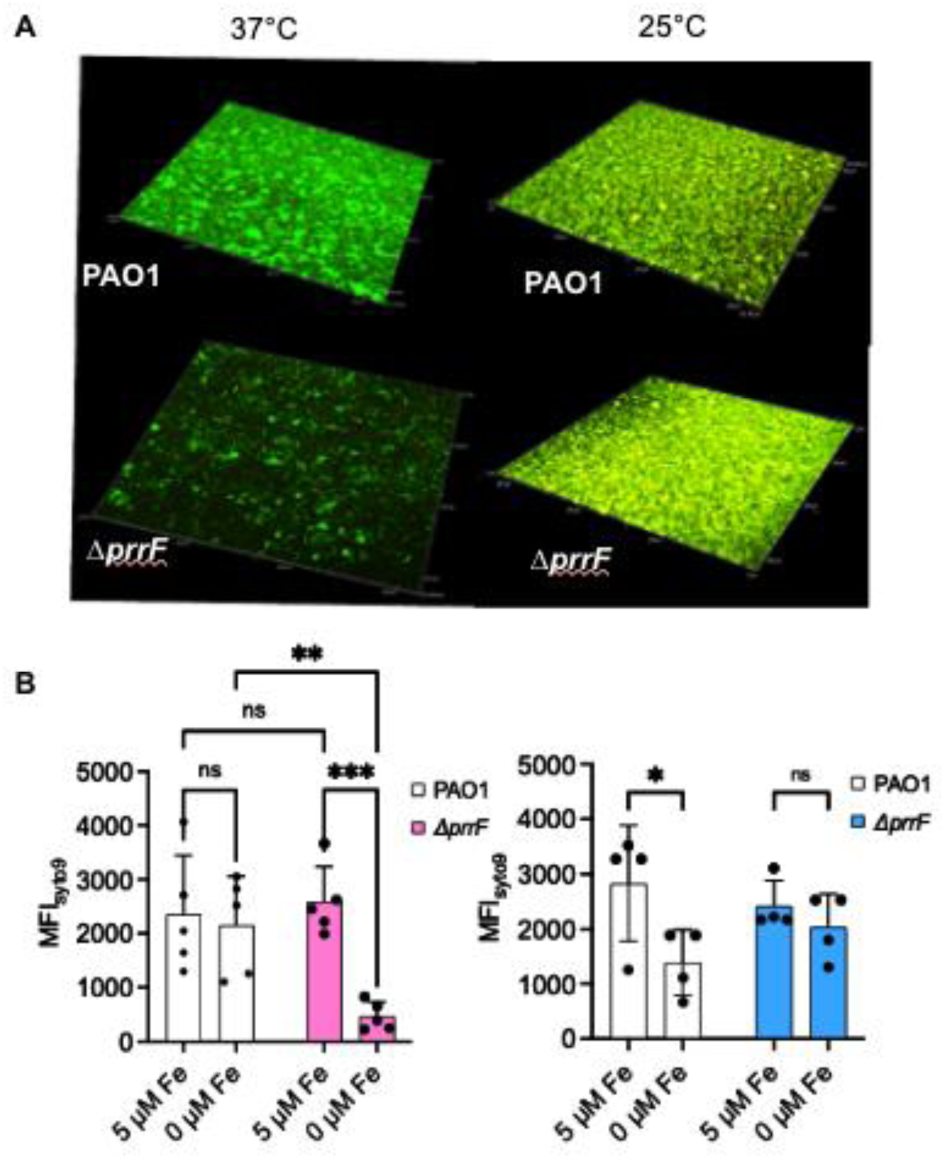
The Δ*prrF* mutant is defective for biofilm formation at 37°C but not 25°C. **A.** Representative confocal microscopy images of the WT and Δ*prrF* strains grown in flow-cell biofilms at either 37°C or 25°C as described in the materials and methods. Live cells are stained with SYTO9 (green) and eDNA with propidium iodide (red). The red fluorescence of PI has been pseudo-colored with orange. **B.** Quantitation of the max fluorescent intensity (MFI) of the indicated conditions from least 5 independent biofilm replicates. Images from all biological replicates are shown in the Supplementary Materials (**Fig. S1-2**).

**Figure 3.**
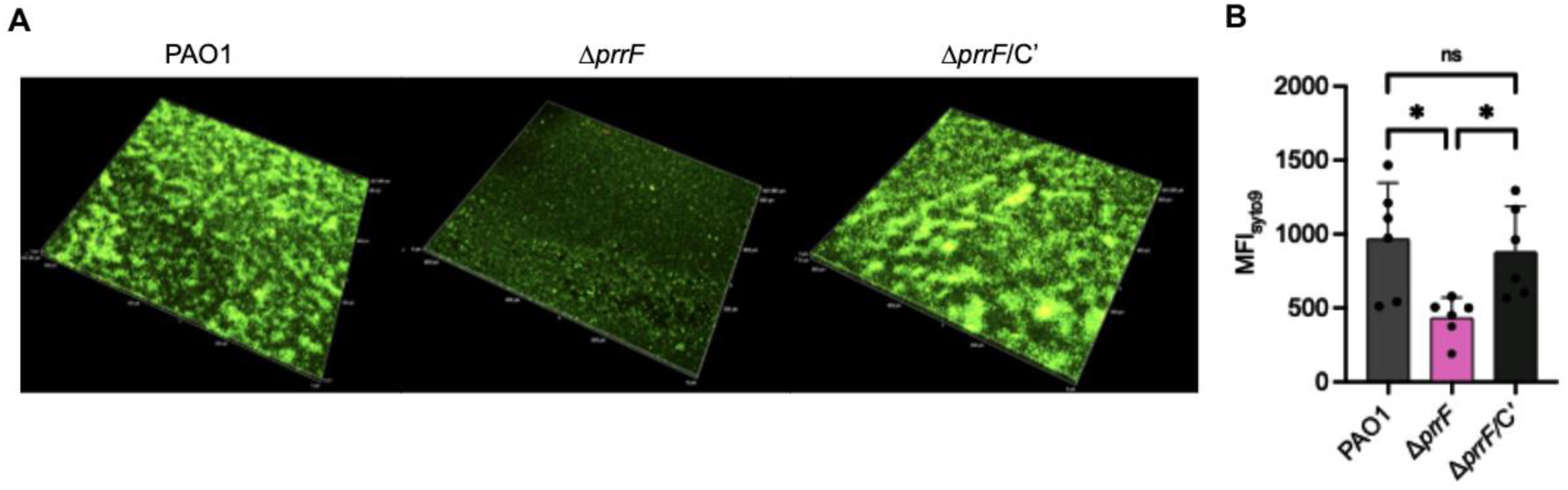
The defect in biofilm formation in Δ*prrF* mutant at 37°C is reversed by complementing with PrrF (Δ*prrF*/PrrFC). **A.** Representative confocal microscopy images of the WT/vector, Δ*prrF*/vector and Δ*prrF*/PrrFC’ strains grown in flow-cell biofilms at 37°C as described in the materials and methods. **B.** Quantitation of the max fluorescent intensity (MFI) of the indicated conditions from least 5 independent biofilm replicates.

### PrrF negatively affects known PrrF-regulated transcripts at 25°C

We next examined whether decreased temperature affected PrrF expression and/or function in planktonic cultures. Real time PCR (qPCR) demonstrated that the PrrF sRNAs are expressed in low iron conditions at 25°C, with PrrF C_T_ levels that are similar to what we observe when PAO1 is grown in low iron media at 37°C (**Fig. 3A-B**). Similar to earlier reports at 37°C, we also observed robust repression of the PrrF sRNAs when PAO1 was grown in medium supplemented with 100 µM FeCl_3_ at 25°C (19, 22, 25, 31, 32). Reporters for two distinct PrrF target mRNAs (PA4880 and *antR*) were strongly de-repressed in the Δ*prrF* mutant in iron-depleted media at both 37°C and 25°C (**Fig. 3B-F**, pink bars at 37°C, blue bars at 25°C), demonstrating PrrF still represses expression of these genes at 25°C. While the PrrF sRNAs may regulate a distinct set of targets at 25°C, these data demonstrate that the PrrF sRNAs are functional and regulate at least some of the same targets at environmental temperature.

### The *prrF* mutant lacks extracellular appendages at body temperature

To examine the potential role of cell surface appendages in the Δ*prrF* mutant biofilm growth defect, we performed transmission electron microscopy of wild type and Δ*prrF* cells grown in a chemically defined medium (CDM) with or without iron supplementation at 37°C and 25°C. Our data shows presence of surface appendages in PAO1 strain which are absent in Δ*prrF* mutant (**Fig. 5**). These surface appendages are also seen in Δ*pilA* mutant indicating these are independent of type IV pili. Notably, these appendages are only present when PAO1 is grown at 37°C under iron-starved conditions. Taken together, our data indicate that PAO1 produces extracellular appendages in PrrF-dependent manner but only at 37°C (**Fig. 5**).

**Figure 4.**
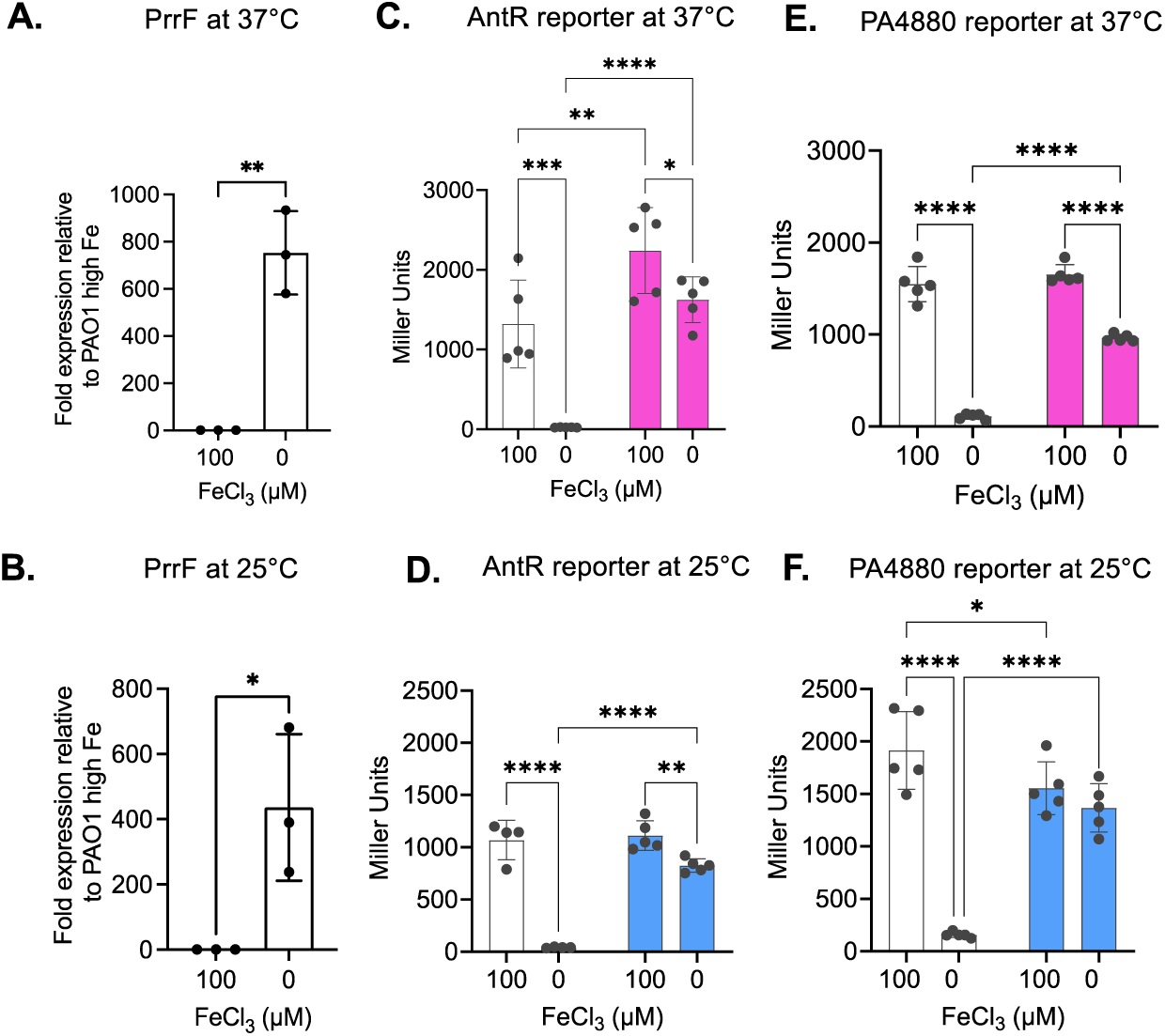
PrrF expression and regulation of two known targets is maintained at 25°C. The indicated strains were inoculated into DTSB, with or without supplementation of 100 μM of FeCl_3_, and incubated for 18 h at either 25°C or 37°C. Cultures were incubated under shaking conditions at 250 rpm. (**A and B**) RNA was isolated and used for RT-PCR as indicated in the Materials and Methods. qPCR results represent means ± SD of 3 biological replicates, with 3 technical replicates included for each biological replicate. (**C-F**) Reporter assays were performed by using the indicated strains containing the *P_antR+UTR_-LacZ^-SD^* and *P_pa4880+UTR_-LacZ^-SD^* reporter fusions. Cultures were subjected to β-galactosidase activity assay as described in the Materials and Methods. Reporter assay data represents mean ± SD of 5 biological replicates. Individual data points for all graphs are represented by a solid black circle. Asterisks indicate significant difference between indicated horizontal bars. *P<0.05, **P <0.005 and ***P <0.001, ****P < 0.0001 by unpaired t-tests for the qPCR data and two-way ANOVA with Fisher’s LSD test for multiple comparisons for the reporter assays.

**Figure 5.**
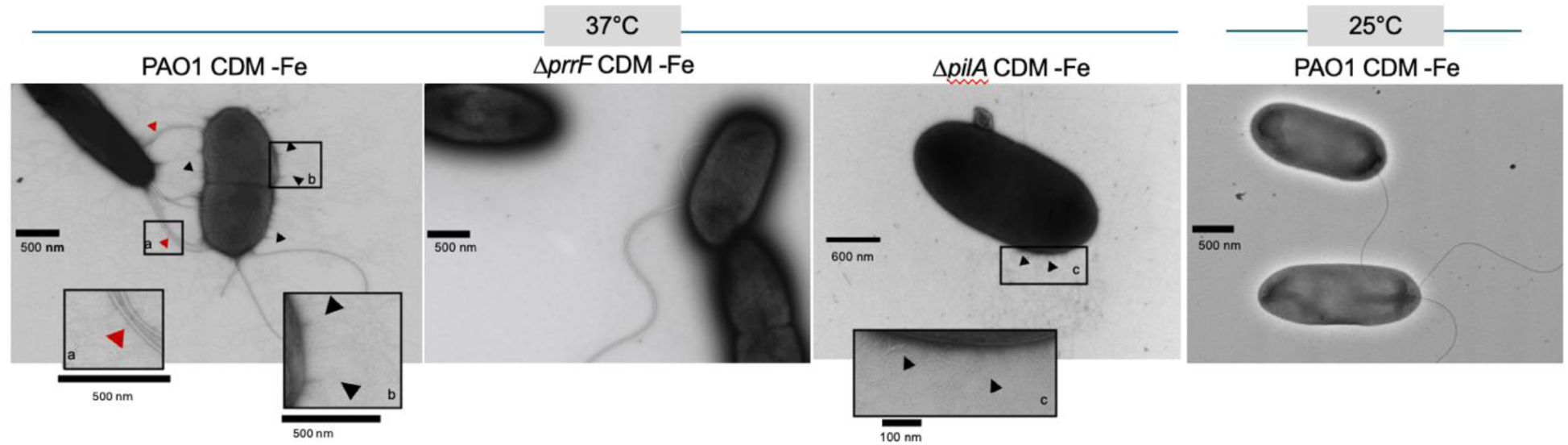
Extracellular appendages are produced in iron starved conditions in a PrrF-dependent manner only at 37 ⁰C. The strains were grown in CDM with 0.1uM Cu, 0.1uM Ni, 1mM Ca, 0.3uM Mn, 6uM Zn for 18 hours at 37 ⁰C shaking at 250 rpm. The samples were loaded onto formvar/carbon-coated grids and stained with 0.25% uranyl acetate. Shown are representative images from at least 3 biological replicates.

## DISCUSSION

This study demonstrates, for the first time, the importance of the PrrF sRNAs and temperature in *P. aeruginosa* biofilm formation. While the specific mechanism underlying this finding remains unknown, this is a critical discovery considering that many flow-cell biofilm studies with *P. aeruginosa*, particularly those regarding iron, have been conducted at 25⁰C (21). We further show that PAO1 produces an unknown surface appendage at 37⁰C, and that this appendage is lacking in both the Δ*prrF* mutant grown at 37⁰C as well as wild type PAO1 grown at 25⁰C. These appendages were not affected by deletion of *pilA*, encoding the major type IV pilus protein that is required for initiation and development of *P. aeruginosa* biofilms (33). These appendages bare striking resemblance to Cup fimbriae that are observed in a mutant lacking both type IV pili and flagella and overproducing the RocS1 sensor kinase (34). While not conclusive, the correlation of these appendages with PrrF-dependent biofilm formation provides an intriguing hypothesis: that Cup fimbriae are key drivers of 37⁰C biofilm formation in low iron conditions. PAO1 encodes four distinct *cup* loci, each of which provide all the genetic components needed to form the Cup fimbriae (34). Current studies in our group are aimed at whether these appendages are indeed Cup fimbriae and, if so, which loci are responsible for their production in a temperature- and PrrF-dependent manner.

Temperature is a key factor for virulence gene regulation of many pathogens, yet limited studies have addressed how temperature affects global gene regulation in *P. aeruginosa* (**35, 36**). *P. aeruginosa* is the only well-studied pseudomonad that grows at body temperature and can cause infection, suggesting growth at 37°C is a step in its evolution as an opportunistic pathogen. Two independent studies showed that *P. aeruginosa* significantly changes in its transcriptome when grown at 37°C versus 25°C. One of these studies discovered a shift from pyoverdine to pyochelin siderophore gene expression at 25°C compared to 37°C, suggesting Fe homeostasis plays a key role in the transition from environmental to body temperature (**35, 36**). This begs the question whether the same observed requirement for pyoverdine but not pyochelin will also be seen in 37°C biofilm formation by *P. aeruginosa*.

RNA thermometers have become increasingly appreciated in the role of temperature-dependent gene expression changes. These elements are often found in the 5’ untranslated regions (UTRs) of bacterial virulence genes and melt upon increased temperature, thereby altering access of RNA polymerase to the translational start site (37–39). To our knowledge, three RNA thermometers have been identified in *P. aeruginosa* mRNAs: these encode the RhlI and LasI quorum sensing synthases and PtxS which activates transcription of Exotoxin A (40, 41). Notably, the *toxA* gene encoding exotoxin A is also induced by iron starvation, providing yet another link between temperature and iron regulation. These and other mechanisms likely alter the ability of certain genes to be expressed at either environmental or body temperature, potentially affecting interactions between PrrF and its target mRNAs. The role of RNA structure in PrrF-mediated biofilm formation is therefore a high priority for future studies regarding temperature dependent changes in *P. aeruginosa* biofilm physiology.

It is important to note that findings from room temperature biofilm studies remain clinically significant even if they are distinct from what is observed at body temperature. *P. aeruginosa* is commonly acquired in hospital settings and has been identified in environmental reservoirs such as sink faucets (42) where it likely grows in biofilms. *P. aeruginosa* must therefore adapt to increased temperature to establish infection in hospital settings. A recent study from Alan Hauser’s group further showed that *Pa* can be shed back into the environment via biofilm “blooms” in the murine gall bladder, leading to intestinal colonization, fecal excretion, and transmission to uninfected cage-mates (43), reminiscent of how gastrointestinal pathogens including *Salmonella* are transmitted during human infection (44). While this finding was restricted to mice, several studies suggest that *P. aeruginosa* can colonize the human gall bladder and intestines, potentially serving as a reservoir in clinical settings (45–50). Thus, the idea that clinical *P. aeruginosa* septicemia is a “dead-end” needs to be reconsidered, with temperature and biofilms playing potentially key roles in transmission to and from the host.

In closing, we have demonstrated that the iron-responsive PrrF sRNAs are important determinants of PAO1 biofilm formation at 37°C, and we predict that additional biofilm determinants of *P. aeruginosa* will vary between environmental and body temperatures. We hope this work provides an increased appreciation for the role of temperature in *P. aeruginosa* physiology and biofilm formationand the impact of temperature-dependent gene regulation in clinical settings.

## MATERIALS AND METHODS

### Bacterial strains and growth conditions

*P. aeruginosa* lab strain PAO1 and the isogenic deletion mutant Δ*prrF* were used for all studies. Strains were routinely grown overnight by streaking from freezer stocks in tryptic soy agar (TSA) (Sigma, St Louis, MO) plates. Three to five isolated colonies were taken from agar plates and inoculated in 2 ml of L broth (LB, Sigma, St Louis, MO) for overnight cultures prior to experiments. Brain heart infusion (BHI) agar was for growth of *Pseudomonas* strains during genetic manipulations. *Escherichia coli* strains for cloning were grown in LB. Antibiotics were added in the following concentration: ampicillin, 100 μg/ml (*E. coli*); tetracycline, 10 or 15 μg/ml (*E. coli*); gentamicin 20 μg/ml (*E. coli*); irgasan 25 μg/ml, carbenicillin, 250 μg/ml (*P. aeruginosa*); tetracycline, 150 μg/ml (*P. aeruginosa*); gentamicin 50 μg/ml (*P. aeruginosa*).

### Generation of PAO1 P*_prrF1_*-*gfp* reporter strain

The promotor region of *prrF1* gene (P*_prrF1_*) was amplified by PCR using primers in **Table S1** and the PAO1 genomic DNA as a template. The vector pMQ37 containing the coding region for green fluorescent protein was linearized by digestion with restriction enzyme HindIII. The linearized vector and P*_prrF1_* PCR product were transformed into yeast (*S. cerevisiae* INVSc1) where the P*_prrF1_* PCR product was incorporated into the pMQ37 vector via homologous recombination. The pMQ37-P*_prrF1_* plasmid was isolated from yeast culture and transformed into *E. coli* strain SM10λ. The pMQ37-P*_prrF1_* plasmid was isolated and digested by restriction enzymes BamHI and EcoRI to clone them into the reporter plasmid mini-CTX-1 to generate mini-CTX-P*_prrF1_*-GFP plasmid. The vector was digested by the same restriction enzymes used for the promoter fusion fragment. The mini-CTX-P*_prrF1_*-GFP plasmid was transformed in *E. coli* strain SM10λ. The mini-CTX-P*_prrF1_*-GFP plasmid was isolated and transformed into *P. aeruginosa* PAO1 via electroporation. pFLP plasmid was isolated and transformed into PAO1 mini-CTX-pPrrF1-GFP via electroporation to excise the integrated mini-CTX after the P*_prrF1_*-GFP reporter construct was introduced into PAO1 at the chromosomal *att* site as previously described (51).

### Generation of PAO1 PA4880 reporter strain

The promoter+UTR (untranslated region) of *pa4880* were amplified by PCR using primers mentioned in **Supplementary Table S1** and the PAO1 genomic DNA as a template. The PCR product was cloned in a TA cloning vector PCR2.1 (Invitrogen), and the sequence was confirmed by sequencing. The promoter fragment was then digested by restriction enzymes EcoRI and HindIII to get them cloned into a promoter-less *lacZ* reporter plasmid Mini-CTX-*lacZY*^-SD^ vector which also lacks the *lacZ* Shine-Dalgarno site to generate *P_pa4880+UTR_-LacZ^-SD^*plasmid for transcriptional+translational reporter assays. The vectors were digested by the same restriction enzymes used for the promoter fragment. All the reporter constructs were introduced into PAO1, and Δ*prrF* chromosomes by integrating them at the *att* site as previously described (51).

### Real time PCR

Quantitative PCR (qPCR) was performed using 3 biological replicates of wild type *Pseudomonas aeruginosa* PAO1 and the isogenic deletion mutant Δ*prrF*. The strains were grown for 18 h at 37°C in a previously described amino rich medium, CDM (52) supplemented with or without 100 uM FeCl_3_.RNA was isolated, DNase treated, and cDNA were synthesized using reverse transcription kit (Promega) with 50ng of total RNA as previously mentioned (53). The cDNA was used for performing qPCR with Taqman (Takara). During setup of the qPCR reaction, each of the biological replicates had three technical replicates. Primers and probes were generated using IDT, mentioned in supplementary materials, **Table S1**. 16S rRNA was used as a normalizer. qPCR data analysis was carried out using standard curve equations generated from cDNA having 1:10 serial dilutions of RNA for each individual targets.

### Reporter assays

AntR and PA4880 reporter constructs in PAO1 WT and *ΔprrF* backgrounds were assayed for beta-galactosidase activity. Strains were inoculated to an OD of 0.05 into chemically defined media (CDM) supplemented with 1mM CaCl_2_, 0.1µM CuCl_2_, 0.1µM NiCl_2_, 6µM ZnCl_2_ and 0.3µM MnCl_2_. In addition, 100µM FeCl_3_ was added for high iron cultures only and grown shaking for 18 hours at either 25°C or 37°C. Cells were then harvested by centrifugation, resuspended in KPB buffer, then diluted 1:10 in Z buffer; chloroform and 1% SDS was also added into the reaction mixture. ONPG was then used as a substrate to begin the reaction, which was stopped by addition of 1M Na_2_CO_3_, once sufficient yellow color was observed. The OD_420_ was measured for each sample, and the beta-galactosidase activity was calculated using the miller units formula = (1000 x OD_420_)/ (time x volume x OD_600_).

### Biofilm growth

Strains were grown in LB overnight, then diluted to an optical density at 600 nm (OD_600_) of 0.05 and inoculated into a 3-chamber flow cell (IBI Scientific, Dubuque, Iowa). The chamber was placed upside down for 1 hour to allow for bacterial adherence to the glass coverslip surface then turned right side up, and a peristaltic pump (Ismatec, Wertheim, Germany) was used to pump 1% LB (v/v) through the flow cell at a rate of 3.5 mL/hour at 25°C and 37°C for 72 and 48 hours respectively. The media was supplemented with 1mM CaCl_2_, 0.1µM CuCl_2_, 0.1µM NiCl_2_, 6µM ZnCl_2_ and 0.3µM MnCl_2_. Additionally, 5µM FeSO4 was supplemented for high iron conditions. Biofilms were stained with Hoechst 33342 (live cells) or SYTO9 (live cells), and propidium iodide (PI, extracellular DNA in the biofilm matrix) as indicated for 30 minutes, at which point flow with 1% LB was resumed to remove excess stain. The biofilms were imaged between the inlet and center of chambers as indicated using a Nikon A1 Confocal Microscope (Melville, NY) and captured with NIS Elements software, using a 20X objective.

### Electron microscopy

Bacterial strains were grown overnight in CDM without iron supplementation. The cultures were harvested and washed in CDM. Next, they were diluted 1:10 in CDM. 5 μl of sample were loaded on the glow-discharged, formvar carbon-stabilized grid and let adsorb for 1 min. Excess sample was removed from the grid using filter paper. The grids were then fixed with 4 % glutaraldehyde in 0.12 M sodium cacodylate buffer pH 7.2 for 5 min.

Excess glutaraldehyde was removed by filter paper and the grids were washed 4 times with sterile distilled water and stained with 0.25 % uranyl acetate for 60 s. Excess stain was wicked off the grid with filter paper, and the grids were allowed too air-dry. Samples were imaged with an FEI tecnai T12 (Thermo Fisher) transmission electron microscope at 80 KV with an AMT bottom-mount camera.

